# Coiled-coil domains are sufficient to drive liquid-liquid phase separation of proteins in molecular models

**DOI:** 10.1101/2023.05.31.543124

**Authors:** Dominique A. Ramirez, Loren E. Hough, Michael R. Shirts

## Abstract

Liquid-liquid phase separation (LLPS) is thought to be a main driving force in the formation of membraneless organelles. Examples of such organelles include the centrosome, central spindle, and stress granules. Recently, it has been shown that coiled-coil (CC) proteins, such as the centrosomal proteins pericentrin, spd-5, and centrosomin, might be capable of LLPS. CC domains have physical features that could make them the drivers of LLPS, but it is unknown if they play a direct role in the process. We developed a coarse-grained simulation framework for investigating the LLPS propensity of CC proteins, in which interactions which support LLPS arise solely from CC domains. We show, using this framework, that the physical features of CC domains are sufficient to drive LLPS of proteins. The framework is specifically designed to investigate how the number of CC domains, as well as multimerization state of CC domains, can affect LLPS. We show that small model proteins with as few as two CC domains can phase separate. Increasing the number of CC domains up to four per protein can somewhat increase LLPS propensity. We demonstrate that trimer-forming and tetramer-forming CC domains have a dramatically higher LLPS propensity than dimer-forming coils, which shows that multimerization state has a greater effect on LLPS than the number of CC domains per protein. These data support the hypothesis of CC domains as drivers of protein LLPS, and has implications in future studies to identify the LLPS-driving regions of centrosomal and central spindle proteins.

**STATEMENT OF SIGNIFICANCE:** The liquid-liquid phase separation of coiled-coil proteins has been implicated in the formation of membraneless organelles such as the centrosome and central spindle. Little is known about the features of these proteins that might drive their phase separation. We developed a modeling framework to investigate the potential role of coiled-coil domains in phase separation, and show that these domains are sufficient to drive the phenomenon in simulation. We additionally show the importance of multimerization state on the ability for such proteins to phase separate. This work suggests that coiled-coil domains should be considered for their contribution to protein phase separation.

## INTRODUCTION

Liquid-liquid phase separation (LLPS) is important for the formation of biomolecular condensates. LLPS occurs in cells when macromolecules interact favorably with each other, demix from bulk cytosol, and form a concentrated coexisting phase. This phase can be liquid-like or become more gel-like, loosing its liquid character (1, 2). Here we use LLPS broadly to describe condensates of varying visocoelastic characteristics. LLPS provides a mechanism for cells to localize protein and chemical function (3) without the need for membrane enclosure into aptly named membraneless organelles (3–6). Examples of membraneless organelles that are thought to form via LLPS include P-bodies, nucleoli, stress granules (7, 8), centrosomes (9–11), and the central spindle (12). The formation of condensates *in vitro* is well supported but the role of biologically relevant LLPS, specifically in the biogenesis of membraneless organelles, is actively debated.

There is recent evidence that coiled-coil (CC) proteins have the propensity for LLPS. The formation of the centrosome and central spindle in particular seems to rely on the LLPS of CC proteins. Examples of phase-separating centrosomal proteins include spd-5 (13), centrosomin (14), and pericentrin (15, 16). MAP65 and its mammalian homolog PRC1, both of which help form the central spindle, can undergo LLPS as well (12). Recent studies with spd-5 provide evidence that interactions between its CC domains drive its assembly (17, 18); experiments with pericentrin and centrosomin also suggest an important role of their CC domains in LLPS (14, 16). There are other CC proteins that LLPS, such as the Golgi structural protein GM130 (19), the measles phospho-protein (20), transcription factors FLL2 (21) and TAZ (22), the endocytic trafficking protein Ede1 (23), a protein encoded by retrotransposable element LINE-1 ORF1 (24), the RNA-binding protein Whi3 (25), and a carboxysome-positioning protein McdB (26). These studies support a narrative that CC domains are involved in LLPS, but it is unknown how they might be contributing and if they themselves are sufficient to drive the process. We are motivated by these examples of phase-separating CC proteins and in this paper we answer the question: Could coiled-coil domains themselves be sufficient to drive the LLPS of proteins?

LLPS is typically driven by multivalency (27–29), which is the ability of molecules to form many attractive interactions with other, similar molecules. A useful conceptual framework for LLPS driven by multivalent interactions is the stickers-and-spacers model of associative polymers (27, 30). In this framework, stickers are specific macromolecular components (e.g. residues, domains, or globular patches on proteins) which interact associatively, whereas spacers are the remainder of the macromolecule that tether the stickers together. This framework applies equally well to intrinsically disordered proteins (IDPs) with stickers being individual residues, as it does for proteins where folded domains are stickers and the spacers are flexible linkers tethering the domains together. Multivalency thus increases as the number of stickers in a protein increases. There are a few examples which show the effect of folded-domain-stickers on protein LLPS, such as the synthetic SH3-PRM system introduced by Li et al. (31) and ubiquitin shuttle proteins p62 and UBQLN2 (32, 33). In the rest of the paper, we use the term *polymeric multivalency* to refer to the number of stickers in a single protein chain.

Multimerization of individual stickers is a plausible, but not well explored, multivalency mechanism in proteins. Most discussion of multivalency in proteins has focused on dimers, rather than interactions of larger valency. This is likely because sticker interactions are often assumed to be 1:1, meaning the entirety of one sticker interacts only with the entirety of one other sticker. This is an assumption that comes from the simplifications used in polymer theory models e.g. theory of associative polymers (27, 30). Higher orders of multimers likely occurs naturally between residue-level stickers, however. Charged (34) and polar amino acids can engage in multiple interactions simultaneously, for example in the coordination of metal ions (35, 36). Multivalent interactions for residue-level stickers can be an important factor in protein LLPS (37, 38). Multimerization between protein domains can also occur naturally to increase LLPS propensity (33), and this requires treating entire protein domains as individual stickers. There are few recent studies which have explored multimerization and its effects on LLPS. Carter et al. (39) showed that mutants of TDP-43 containing a tetramerization domain have higher LLPS propensity than wild-type TDP-43 with its dimerization domain. Additional examples of LLPS affected by multimerization have been reported (summarized by Mohanty et al. (33)), but beyond these few the role that multimerization plays in affecting multivalency has not been well explored. In this paper we use the term *multimeric multivalency* to refer to the number of partners that an individual sticker is capable of interacting with at once. We use the familiar terms dimer, trimer, tetramer, etc., to describe specific varieties of multimers.

CC proteins might utilize both polymeric and multimeric multivalency mechanisms to drive LLPS through their CC domains. If we apply the stickers-and-spacers framework to CC proteins and treat the CC domains as the stickers, we see that these proteins harness both polymeric and multimeric multivalency. CC proteins can show varying degrees of polymeric multivalency dependent on the number of CC domains in the protein. CC domains also have intrinsic multimeric multivalency because of their ability to form dimers (40), trimers, tetramers, and larger multimers (41). Proteins containing many CC domains could realize LLPS through a combination of polymeric and multimeric multivalence mechanisms; the extent to which this may happen is not understood.

We hypothesize that CC domains are sufficient to drive the LLPS of proteins and that this phenomena can be tuned by both their polymeric and multimeric multivalency. We generated a novel modeling framework to test our hypothesis by coarse-grained molecular simulation (Fig. 1). We use this framework to build CC proteins where CC domains are tethered to other CC domains with disordered, inert linker regions (Fig. 1A). This framework allows us to specify the polymeric multivalency of CC proteins by changing how many CC domains are in a given protein chain (Fig. 1B), as well as the multimeric multivalency of each CC domain (Fig. 1C). For the purposes of this paper, we restrict our attention to dimer-, trimer-, and tetramer-type multimers. We use this framework to focus on general physical properties of phase separation of CC proteins when treated as associative polymers.

**Figure 1:**
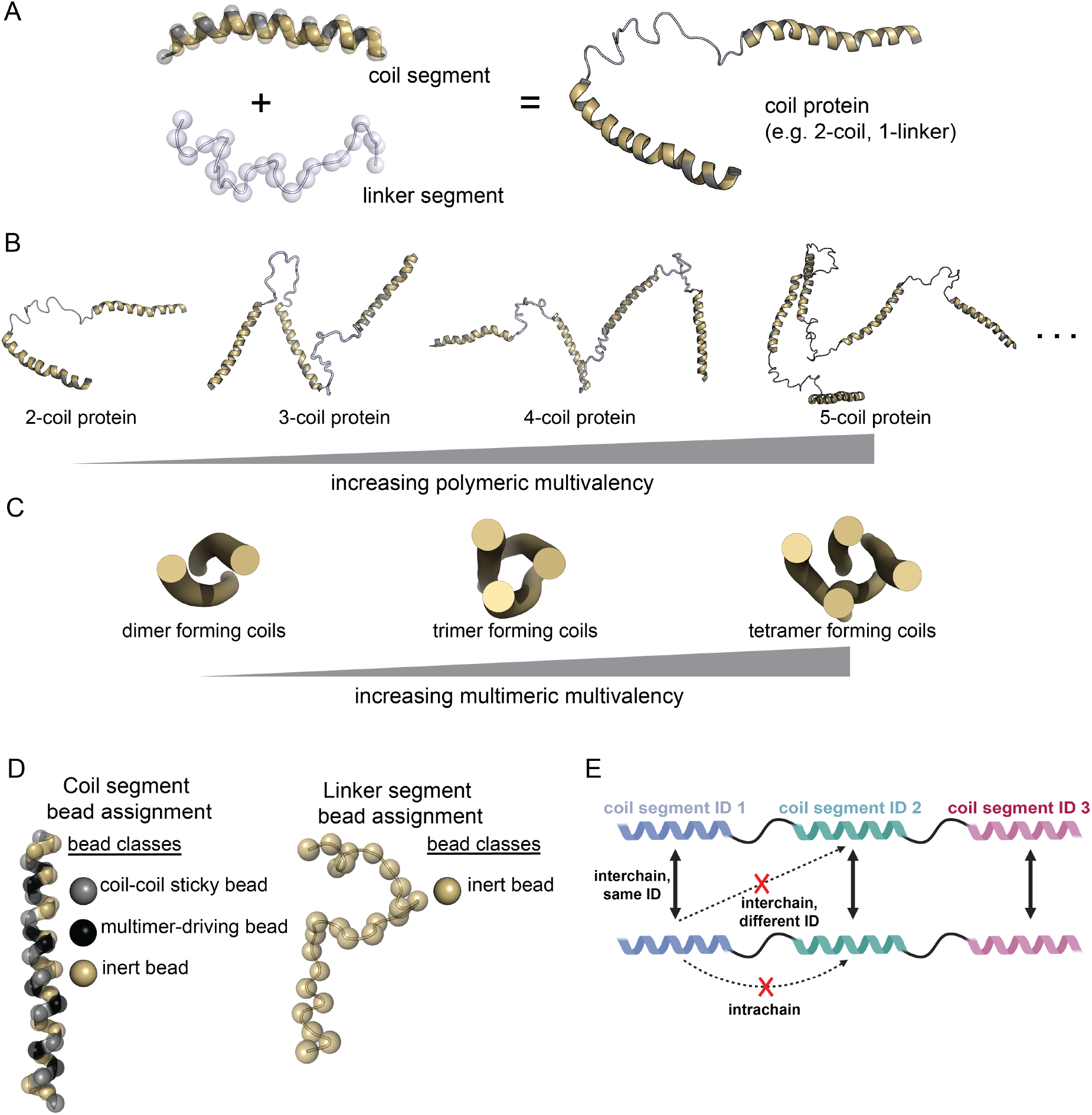
The CC LLPS framework can be used to make CC proteins with varying polymeric and multimeric multivalency. (A) Coil segments and linker segments can be combined in any number to make a coil protein, e.g. a 2-coil-1-linker protein. Haloed spheres represent the CG beads, and are removed in later visualizations for clarity. Helices are the coil segments and the silver portions are the coil-coil sticky beads. Light grey tubes are the linker segments. (B) The polymeric multivalency of a CC protein can be modified by changing the number of coil segments in the protein. Only two through five coil segments are shown, but there could be an arbitrarily large number of coil segments per protein (represented by ellipsis). (C) The multimeric multivalency of a protein can be modified by changing the type of multimer that a coil segment is favored to form. (D) Diagram showing how the different bead classes are organized within each of the segments. (E) Schematic representation of the *maximally specific interaction scheme* used in this study, where only interchain interactions between the same type of coil segment is permitted, indicated by a solid black line. Schematic made using BioRender. All molecular visualizations were produced using open-source PyMOL v.2.5.0.

## METHODS

### Software

All molecular dynamics (MD) simulations were performed using unmodified GROMACS (42) version 2022.1. GROMACS was compiled with GCC version 11.2.0 and Open MPI version 4.1.1 for parallel processing on the Alpine high performance computing resource at the University of Colorado Boulder, and with GCC version 10.2.0 and Open MPI version 4.0.5 for Bridges-2. Non-MPI enabled versions of GROMACS 2022.1 were used for small scale simulations, and some data analysis and processing. These versions were compiled on a Ubuntu machine (22.04.2 LTS) with GCC 11.2.0 and MacOS 12.6 with AppleClang 14.0.0. Additional analysis code was written in Python 3.9.

## Code availability

Files from this study, including molecular dynamics parameter files, relevant topologies, analysis scripts, etc. are available on the GitHub repository associated with this manuscript: https://github.com/dora1300/cc_llps_framework.

### Coarse-grained framework to study coiled-coil driven LLPS

We developed a bespoke, coarse-grained (CG) molecular dynamics framework to study CC LLPS, inspired by other types of CG models of CC proteins (43, 44). Computational methods have been critical in the understanding of protein LLPS (28), but prior to this study there were no tools specifically designed for studying CC-driven LLPS. We designed our framework to be easy to use and readily modifiable so it can represent a variety of different CC proteins of arbitrary size (Fig. 1B), with coils of any desired multimerization state (Fig. 1C), and of defined specificity between coil segments (Fig. 1E).

We define CC proteins as linear combinations of *coil segments* (stickers) and *linker segments* (spacers) (Fig. 1A), which is inspired by the organization of centrosomal proteins (18, 45). Coil segments are regions of the protein which represent CC domains, and linker segments are the tethers between coil segments. We specifically chose this language to avoid confusion with terms like ‘domains’ which can have multiple meanings. This organization allows us to treat CC proteins as associative polymers, where each coil segment acts as a single sticker and each linker segment as a spacer. We implemented the framework with the following objectives and simplifying assumptions in mind: (a) coil segments are helical, and linkers are disordered, (b) protein–protein interactions only occur through coil segments, (c) coil segments retain a heptad-repeat like CG bead organization so that coils interact in a similar geometry to real CC domains, and (d) linkers interact equally between solvent and protein components.

Our framework uses C-α coarse-graining, which represents every amino acid as a single CG bead. These beads are centered at the same coordinates as the C-α carbons of amino acids in a corresponding atomistic representation of a protein. The C-α CG procedure maps traditional ϕ,ψ angles into coarse-grained space (46). The corresponding CG pseudo-bonds and pseudo-torsions allow us to reproduce correct secondary structure geometries in the CG structures. A CG representation thus has significantly fewer beads than an atomistic representation (on average, 19 atoms are replaced by one bead), which greatly increases computational efficiency in molecular simulation, as well as increasing the time step allowed by increasing the masses of the smallest particle. Additional information about the framework’s structural features and interaction parameters (along with details about each bead type used, e.g. Fig. 1D) are provided in Supporting Material, sections S.I and S.II respectively.

### Common molecular dynamics parameters

We used a consistent set of MD run parameters throughout the study, and deviations from these parameters are noted in the respective methods. Energy minimization was performed using steepest descent to a force tolerance of < 50 kJ/mol/nm and a step size of 2 pm. Equilibration and production simulations were done with Langevin dynamics using the sd integrator with a friction coefficient of 0.2 ps^-1^. The center of mass translational velocity was removed every 10 steps. Lennard-Jones potentials were shifted and cut-off at 1.1 nm, consistent with other coarse-grained force fields (47). The verlet-buffer-tolerance, a GROMACS-specific setting for Verlet list updates, was set to 1× 10^−7^. Setting this parameter correctly is important for reproducibility because using default larger numbers for verlet-buffer-tolerance results in simulation instability, due apparently to density inhomogeneity in phase-separated simulation boxes. Random seeds for each simulation were generated pseudo-randomly by GROMACS. Periodic boundary conditions were applied in all three dimensions. We used a single box size for slab simulations to maintain consistent configurational entropy between experiments. The final box size for slab simulations is 25×25×150 nm, which is similar to previous studies (7).

### Temperature replica exchange MD to assess dynamics of coarse-grained CC dimer

We verified that simulated CG CC dimers reproduce similar structure to atomistic simulations of a CC dimer. Thomas et al. (48) performed 100 ns atomistic temperature replica-exchange (REMD) of two, 32-amino acid long coiled-coils starting in a dimer configuration and published the backbone RMSD of each replica. We reconstructed a mean RMSD distribution of their replicas to serve as our reference for a stable, CC dimer (Fig. S2A). We performed simulations of a CG CC dimer in our framework and compared our backbone RMSD distribution to the reference. Specific details are provided in Supporting Material, section S.V. Figure S2B-C shows the backbone RMSD distributions across different coil-coil interaction strengths, at 293 (Fig. S2B) and 310 (Fig. S2C) K. We generated Gaussian kernel density estimates for all of the RMSD distributions and determined the similarity between our CG RMSD data to the atomistic reference distribution by calculating the Kullback-Leibler (KL) divergence (Table S3). The lowest KL divergence, at both temperatures, was from the 5.5 kJ/mol simulations (298 K, 0.90 nats; 310 K, 1.01 nats). Thus, 5.5 kJ/mol is chosen as the optimal coil-coil sticky interaction strength (for dimer-forming coils) for our framework. This validation demonstrates that our CG framework can reproduce a similar structure and equilibrium folding propensity as an atomistic coiled-coil dimer.

### Single molecule MD of intrinsically disordered proteins to parameterize linker segments

We fine tuned the linker segment parameters by systematically weakening the magnitude of the linker structural parameter force constants starting from the coil segment values, and determined the optimal force constants by comparing simulated *R*_*g*_ of 23 IDPs against experimentally determined *R*_*g*_ (details in Supporting Material S.VI). The interaction parameters of the inert beads was left unaltered in all cases. Figure S3 shows a parity plot comparing simulated to experimental *R*_*g*_ for the 23 tested proteins, with structural parameter force constants 100 × weaker than coil segment (i.e. helix-forming) parameters. This level of weakening produced the best correlation between simulated and experimental data. The Pearson correlation coefficient between simulated and experimental data is 0.753. The slope of the simulated versus experimental data is 0.814 and is qualitatively close to the parity line. We judge that this qualitative agreement to real IDPs is sufficient for this study, even in the absence of amino acid-specific chemical information and attractive interactions between the inert linker beads.

### MD to assess multimerization capacity of coil segments

We confirmed that we have moderate control over the multimerization of coil segments in the CC LLPS framework. We verified this by simulating several copies of a 3-coil-2-linker protein with each of the multimer-driving bead types and quantified the multimer species present in the simulations (details provided in Supporting Material section S.VII). Simulations with the dimer multimerization beads (Fig. S4A) show that dimers are the dominant multimer present at about 60% composition over simulation time. Simulations with the trimer multimerization beads (Fig. S4B) produce a mixture of multimers with the dominant multimer species being the trimer at 46% of composition averaged over time. Tetramer multimerization beads result in a structure that still favors trimers but can form tetramers (Fig. S4C), with the trimer and tetramer multimer species comprising 38% and 26% of composition over time, respectively, at a coil sticky interaction strength of 4 kJ/mol. Despite this structure having more trimer than tetramer species, we will refer to it as the tetramer-forming model as it is the only one that forms an appreciable amount of tetramers. Attempts to shift the multimer population towards the tetramer species by changing the coil sticky interaction strength either ablated multimer formation (3 kJ/mol) or resulted in pentamers appearing in the simulation (5 kJ/mol), suggesting that 4 kJ/mol is near optimal strength for tetramer-forming beads (Fig. S4C) in our model framework. The coil sticky interaction parameter (i.e. the effective pair interaction strength parameter) for both the trimer- and tetramer-forming models is weak relative to the dimer-forming model to prevent the formation of aberrantly large multimers. Increasing the interaction strength for all types of multimer-driving bead interactions resulted in multimers larger than desired. The lack of full specificity in the trimer- and tetramer-forming coils might be a fundamental limitation of using a C-α coarse-graining approach, but this remains to be tested.

### Slab simulations for phase coexistence

We used the slab simulation (7, 49, 50) protocol to directly simulate phase coexistence of CC protein models. This procedure involves the following steps: generating configurations of each protein of interest, placing copies of those configurations in a simulation box, compressing the box in one dimension in the NPT ensemble to generate the slab, expanding the box in the same compressed dimension, equilibrating the slab to the desired temperature in the NVT ensemble, and, finally, production simulation in the NVT ensemble. We note that during the packing of the simulation box, copies of proteins are placed randomly throughout, but we fix the total number of protein copies so that all simulations have the same total number of coil segments. This allows us to make comparisons between simulations of different proteins.

#### Single molecule simulations to generate starting configurations of CC proteins

We generated proteins of interest using the `PeptideBuilder` strategy (Supporting Material section S.VIII) and then used MD to equilibrate the structure of each protein and prepare it for slab simulation. A single copy of each protein is placed into a simulation box large enough to accommodate the initial structure. Single molecules were energy minimized and then equilibrated in the NVT ensemble for 500 ps with a time step of 25 fs. NVT equilibrations were used so that the linker segments could relax out of the initial helical configuration and closer to the defined equilibrium angles. At this point, for each protein structure we generated a unique single molecule simulation for every temperature that would be used in the forthcoming slab simulations, i.e. 253, 273, 293, and 313 K. Production MD simulations in the NVT ensemble were performed on all equilibrated structures for 5 μs with a time step of 25 fs at their corresponding temperatures. Only 1 replicate simulation was used for each single molecule simulation at each temperature.

#### Selecting single molecule configurations for slab simulation

We begin the slab protocol by randomly packing copies of our proteins of interest in a simulation box. The single molecule simulations served as the pool of configurations from which we pack the starting box. We standardized the total number of coil segments in a simulation box to 450 coils for all protein models. Simulations with 4-coils per protein had a total of 448 coils in each box. The following protein copy numbers were used to achieve the desired number of coil segments: 225 copies of 2-coil proteins; 150 copies of 3-coil proteins; and 112 copies of 4-coil proteins. We mitigated configurational bias by randomly choosing five configurations from the equilibrated portion of the single molecule simulations (equilibrated state began by 100 ns), using random numbers generated from Random.org. Each of the 5 configurations from each protein was copied so that the total coil density in the simulation would reach the desired number (450). We prepared three replicate simulations at every desired temperature, with different random numbers for each replicate, using only the corresponding single molecule trajectories to provide the structures. packmol v20.11.1 (51) was used to pack all the initial slab boxes, with a tolerance of 1.0 nm between every protein copy, into a box size of 25×25×100 nm to ensure all copies would fit inside. We generated initial slab simulation boxes for every protein of interest, at every desired temperature, in triplicate.

#### Slab simulations

Packed boxes were energy minimized and then equilibrated in the NPT ensemble for 200 ns with a time step of 20 fs at 150 K. The goal of this step is to create the actual protein slab to assess phase coexistence. Semi-isotropic pressure coupling was applied only in the z-dimension of the box. The Parrinello-Rahman barostat was used with a reference pressure of 1 bar, compressibility of 3 × 10^−4^ bar^−1^, and a time constant of 5 ps for the coupling. Simulation boxes are compressed to approximately 10-20% of the starting z-axis length at the end of NPT equilibration. Proteins were “unwrapped” prior to expanding the box to prevent aberrant expansion of individual proteins. The simulation box was edited such that the z-dimension became the final length of 150 nm, and the protein slab was kept centered in the simulation box during expansion. Slabs in expanded boxes were then equilibrated in the NVT ensemble for 200 ns with a time step of 20 fs, at the desired temperatures. We then performed production MD simulations in the NVT ensemble for each temperature-equilibrated slab for a total of 20 μs with a time step of 25 fs, with the GROMACS parameter rdd set to 1.6 nm (important for domain decomposition). We found that long simulation times for our proteins are necessary when compared to other studies with the slab method (7, 50, 52), to ensure that our systems reach equilibrium. Equilibrium was assessed by monitoring the density of both the center of the box, and the opposite ends of the box combined. We considered the simulation/slab to be equilibrated when the density of both these regions was stable. Only the equilibrated portions of trajectories were used for analysis. Almost half of all simulations equilibrated within 2.5–5 μs, but some simulations, particularly the 4-coil containing proteins and tetramer-forming coils, took up to 10 μs to equilibrate.

### Density profile analysis and generation of binodals

We performed density analysis of each equilibrated trajectory using the gmx_density module in GROMACS. Density (reported as number density / nm^3^) was calculated in 2 nm slices along the z-axis of the simulation box, and the average density in each 2 nm slice for a trajectory is output. Density profiles (which are reported along the z-axis) for a given protein at a given temperature were averaged across three replicate simulations and are plotted as mean (solid line) ± standard deviation (shaded region). These data are used in determining LLPS, in combination with molecular cluster analysis (described below). We also used the density output to calculate binodals. For a single trajectory, number densities in the center of a slab corresponding to 70–80 nm in the z-dimension was averaged to describe the dense region. Number densities at the edges of the box not corresponding to a slab, ranging from 1–35 and 115–150 nm in the z-dimension, were averaged to represent the dilute region. Dense and dilute regions across triplicate simulations were averaged and plotted along with standard deviation in binodals.

### Molecular cluster analysis

Molecular cluster size distributions were determined from every 10 ns of equilibrated trajectories using the gmx_clustsize module in GROMACS. Cluster sizes for whole molecules were calculated using a distance cut-off of 0.9 nm between interacting pairs. The output cluster size populations were normalized to probabilities and averaged for each temperature replicate for each protein of interest, and are reported as mean ± standard deviation over three replicate simulations. Molecular cluster analysis is used to help determine LLPS in slab simulations, in combination with density profile analysis. Both a molecular cluster containing nearly all of the proteins in a simulation and the simultaneous presence of a density transition in the *z*-direction are markers of LLPS.

### Mean squared displacement and diffusion analysis

The average mean squared displacement (MSD) for each type of protein was calculated using the gmx_msd module in GROMACS. The MSD of individual protein molecules was calculated from each molecule’s center of mass. Analysis occurred every 10 ns from the final 10 μs of each trajectory. Molecules were made whole, and periodic boundary conditions were accounted for in all three dimensions. The configuration at time 10 μs was used as the reference. Individual protein MSD data were then averaged for each replicate. Effective diffusion coefficients at all temperatures for proteins that form LLPS droplets were estimated from short-time MSD data (from 1 to 5 (μs) lag-times) via bootstrapping. 450 data points were sampled with replacement randomly from any of the three replicate MSD data sets for any given protein. A line was fit to each set of 450 randomly sampled points, and the diffusion coefficient was calculated by dividing the estimated slope by six. The averaged effective diffusion coefficient and standard deviation was calculated from 5000 bootstrap cycles. Due to long-timescale kinetic trapping in simulations with LLPS, we do not expect MSD to be linear through all lag-times. Thus the effective diffusion coefficients are likely high compared to the true diffusion coefficients for each system. Effective diffusion coefficients were not corrected for box size, as only the qualitative relative differences in diffusion constants are of interest.

## RESULTS

### Dimer-forming coil segments can drive LLPS of CC proteins

We hypothesized that CC proteins would readily form LLPS droplets in simulation due to physical features of CC domains that could enable both polymeric and multimeric multivalency mechanisms. We developed a custom simulation framework to study the phase separation of CC proteins, and observed LLPS over a range of polymer multivalency, multimer multivalency, and temperatures.

We first investigated the minimum number of dimer-forming coil segments that are needed in a protein to see LLPS. We tested three coarse-grained CC proteins — 2-coil-1-linker, 3-coil-2-linker, and 4-coil-3-linker (Fig. 1B, first three proteins shown), all with dimer-forming coils — for LLPS propensity using the slab method. A LLPS droplet is determined quantitatively from density profile and molecular cluster analyses and exists when there is both a sharp density transition and the presence of a molecular cluster whose size is close to the number of proteins in the simulation. Averaged density profiles and molecular cluster size distributions for all simulated proteins are reported in Figure S5.

All three CC proteins formed LLPS at 253 K, the lowest of the temperatures that we simulated (reflected in representative snapshots in Fig. 2 and in phase diagrams in Fig. S8A). This demonstrates that proteins with as few as two coil segments can LLPS at low temperatures. At higher temperatures (T ≥ 293 K), LLPS is not observed and the proteins are dilute throughout the system. The 3-coil and 4-coil proteins also formed LLPS droplets at 273 K, unlike the 2-coil protein. Increases in polymeric multivalency is known to increase LLPS propensity (4, 31, 53, 54), and our observations with dimer-forming coils are in line with these reports. Figure 3 shows a binodal plot for the 3-coil dimer-forming protein (Fig. 3A) along with a series of snapshots at each temperature corresponding to the points on the binodal (Fig. 3B). Snapshots of 2-coil (Fig. S6A) and the 4-coil (Fig. S6B) dimer-forming proteins at each temperature are also presented. These combined data (Figs. 3 and S6) highlight that high temperatures destabilize LLPS droplets and result in dispersed proteins in simulation. These data demonstrate that dimer-forming coil segments are sufficient to drive LLPS of proteins, and that increasing polymeric multivalency slightly increases LLPS propensity.

**Figure 2:**
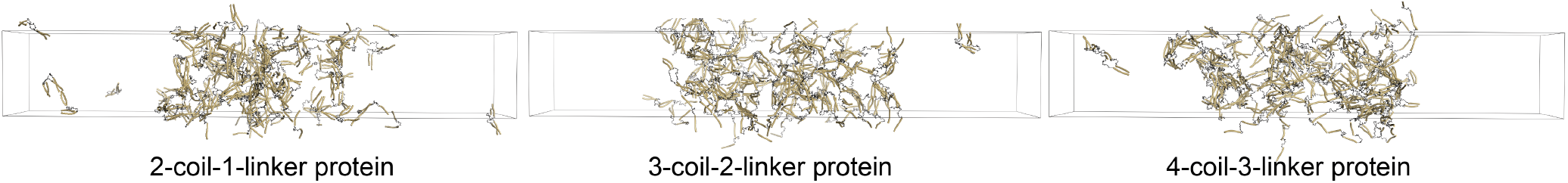
CC proteins with varying number of coil segments all phase separate. Images from slab simulations of 2-coil-1-linker (left), 3-coil-2-linker (middle), and 4-coil-3-linker (right) proteins show LLPS is supported at varying polymeric multivalency for dimer-forming coils. Snapshots are from simulations at 253 K and 20 μs. Proteins are made whole for visualization, but would wrap through the box boundaries during actual simulation. Visualizations produced using open-source PyMOL v.2.5.0.

**Figure 3:**
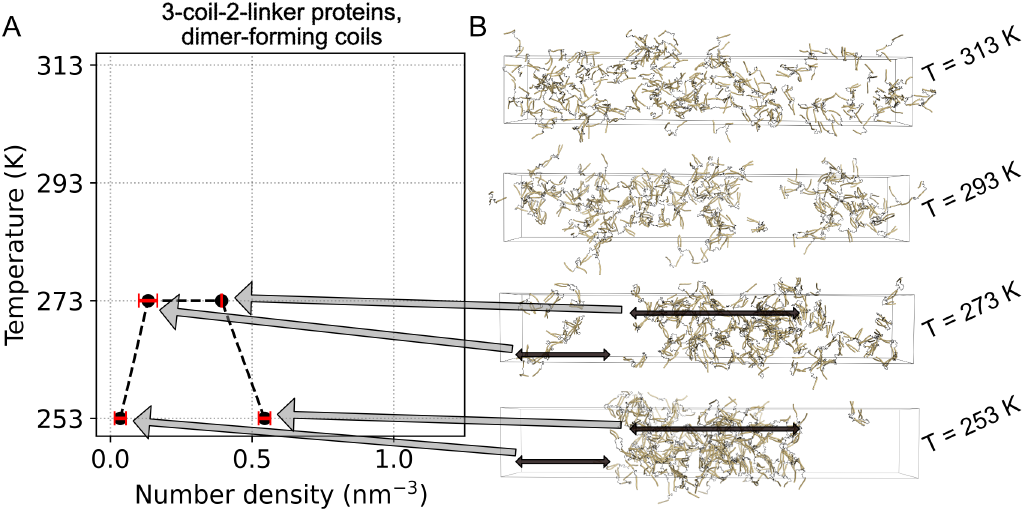
LLPS behavior varies with temperature for dimer-forming coil proteins. (A) Binodal plot for a 3-coil-2-linker protein, which also appears in Fig. 5B. Data points on the left-hand side are for dilute regions, and right-hand side are for dense (droplet) regions. No points are shown at 293 K and above as no phase separation occurs at those temperatures. Data points are presented at mean (circles) ± standard deviation (red bars) from three replicate simulations at each temperature. (B) Snapshots (at 20 μs) from slab simulations from one replicate set for a 3-coil-2-linker protein with dimer-forming coils. Grey arrows between panels corresponds to points on the binodal and associated regions within a simulation box, and black arrows highlight the extent of each region. The snapshot at 253 K is the same as in Figure 2. Proteins are made whole for visualization, but would wrap through the box boundaries during actual simulation. Density profiles for each temperature shown here can also be seen graphically in Figure S5A, left column, middle row. Visualizations produced using open-source PyMOL v.2.5.0.

### Multimerization greatly affects LLPS propensity, whereas polymer multivalency only weakly affects LLPS propensity

We varied the multimeric state of the three proteins (2-coil, 3-coil, and 4-coil proteins) to make trimer-forming and tetramer-forming versions. We ran slab simulations on these proteins and analyzed the trajectories for evidence of LLPS. Density profiles and molecular cluster distributions show an increasing propensity for LLPS as multimer state increases (Fig. S5). We inspected the trajectories and observed a multimer-state dependent slab compaction, as demonstrated in Figure 4 for 2-coil proteins, whereby increasing the multimer state results in a more dense and compact LLPS droplet. This behavior is also demonstrated graphically in Figure 4A which additionally highlights a sharp density transition indicative of LLPS. This compaction behavior was observed for 3-coil and 4-coil proteins as well when inspecting representative snapshots (Fig. S7) and density profiles (Fig. S5A).

**Figure 4:**
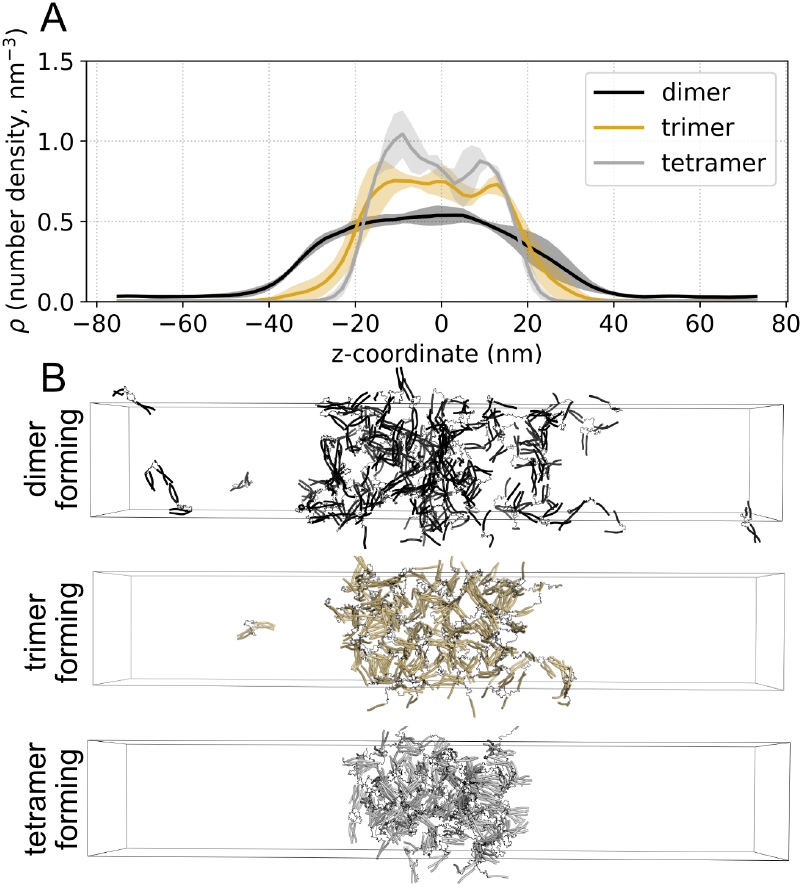
Multimer multivalency is associated with LLPS droplet compaction. (A) Density profiles for dimer-, trimer-, and tetramer-forming 2-coil-1-linker proteins at 253 K. Solid lines represent mean, and shaded regions are standard deviation, from three replicate simulations. (B) Snapshots (at 20 μs) from simulations shown in panel A. Snapshot of dimer-forming protein is same as Figure 2A, but with different coloring of proteins. Proteins wrap through the box boundaries during simulation, but atoms are shifted into neighboring cells to keep proteins visually intact. Colors are coordinated between panels A and B. Visualizations produced using open-source PyMOL v.2.5.0.

We next quantified the impact that polymeric and multimeric multivalency has on the LLPS propensity of our proteins. Our simulations show two results: (1) polymeric multivalency only slightly impacts LLPS of CC proteins, and (2) for a protein with fixed polymeric multivalence, increasing the multimer multivalency dramatically increases LLPS propensity. These results are shown by phase diagrams which compare the phase behavior for all tested proteins (Fig. S8). Increasing polymeric multivalency slightly increased LLPS for dimer-forming coils (Fig. S8A), but we did not see a similar trend for trimer-forming (Fig. S8B) or tetramer-forming (Fig. S8C) coils. We found that as the multimeric multivalency of the system increased, for a fixed polymer multivalence, the LLPS propensity was more robust, i.e. the apparent melting temperature for all proteins increased as the coil multimer-state increased (Fig. S8). The behavior of the 3-coil and 4-coil tetramer forming proteins might differ above 313 K, but additional simulations closer to their melting points are needed.

We constructed binodals from all of the simulations for a qualitative picture of LLPS for each of the proteins. Figure 5 shows the binodals for the (A) 2-coil-1-linker, (B) 3-coil-2-linker, and (C) 4-coil-3-linker proteins, as coil-multimeric state is varied. Data points on the left side of each plot correspond to average densities from the dilute region of the simulation box, and data points on the right side of each plot correspond to average densities from the dense region. We omitted points at temperatures where the dense and dilute regions have overlapping number density, as this represents a melted regime and this data is not included in binodals. These data clearly show that increasing multimeric multivalency strongly increases LLPS propensity and melting temperature of protein droplets. The combination of Figure S8 and Figure 5 demonstrate that polymeric multivalence has a relatively small impact on the LLPS propensity of our CC proteins, whereas multimeric multivalence has a much greater influence.

**Figure 5:**
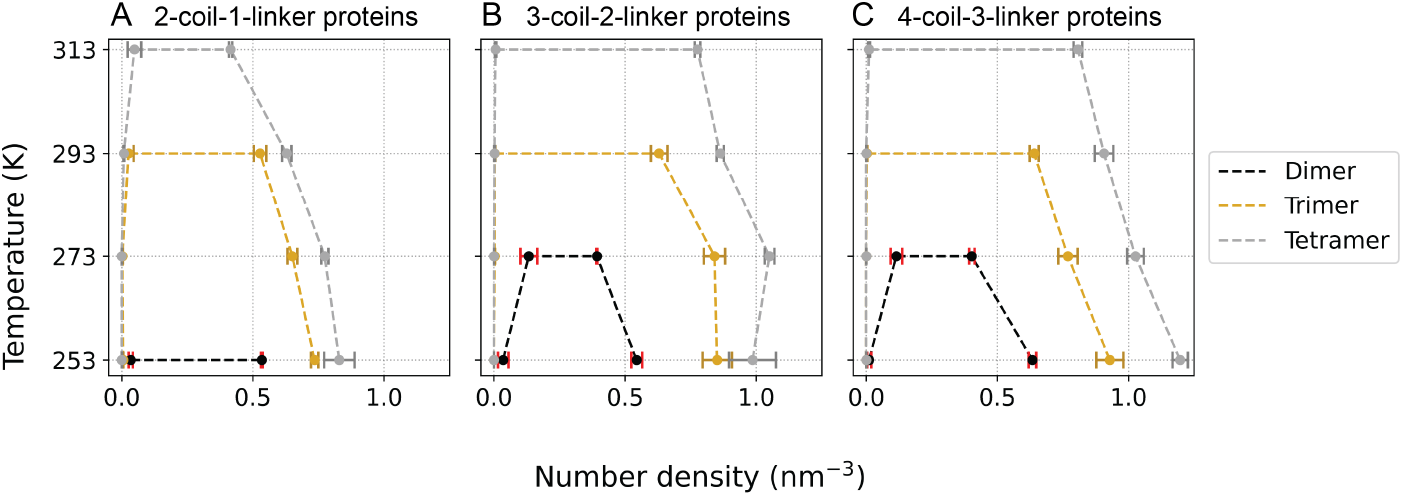
Multimeric multivalency greatly impacts LLPS propensity. Binodal plots for (A) 2-coil-1-linker proteins, (B) 3-coil-2-linker proteins, and (C) 4-coil-3-linker proteins, compare the effect that multimerization has on LLPS propensity. Dimer-, trimer-, and tetramer-forming proteins are represented as black, gold, and silver, respectively. Data points on the left-hand branch of plots are for dilute regions, and on the right-hand branch are for dense regions. Data points are presented as mean (circles) ± standard deviation (bars) from three replicate simulations.

### Protein dynamics within LLPS droplets are affected differently by polymer and multimer multivalencies

Following the multimer-dependent slab compaction observation (Fig. 4 and S7), we characterized the dynamics of proteins within slabs and hypothesized that increasing multimer states reduces the dynamics of individual proteins. We calculated the mean squared displacement (MSD) of individual proteins for all simulations where LLPS occurs (Fig. S8) to estimate the effective diffusion coefficient of proteins within a droplet, with the caveat that proteins in a droplet may be kinetically trapped and thus only partially diffusive on these time scales. Effective diffusion coefficients, estimated by bootstrap on MSD analyses, are plotted in Figure 6 and listed in Table S4. Linear fits from bootstrap analysis used to estimate the diffusion coefficients are shown on MSD plots in Figures S9–S10. Generally, both polymeric and multimeric multivalency reduce protein diffusion, but increasing multimer state does so more drastically than polymeric state (Fig. 6). The effect of multimer and polymer state on diffusion is most clear at 253 K (Fig. 6A), the condition in which all proteins form LLPS. At this temperature, both trimer- and tetramer-forming proteins have dramatically slower effective diffusion coefficients than dimer-forming proteins. The largest difference is between 2-coil proteins of varying multimer states, where dimer-forming coils diffuse ∼ 25 and ∼ 2100 fold faster than trimer- and tetramer-forming coils, respectively. Increasing polymer state also decreases protein diffusion for all proteins at 253 K, but the effect is less evident for trimer- and tetramer-forming proteins due to the strong effect of multimerization on reducing protein dynamics in droplets. Multimer and polymer multivalency-dependent slowing of diffusion is seen at the other temperatures (Fig. 6B–D) but fewer comparisons between proteins can be made due to reduced LLPS propensity at higher temperatures.

**Figure 6:**
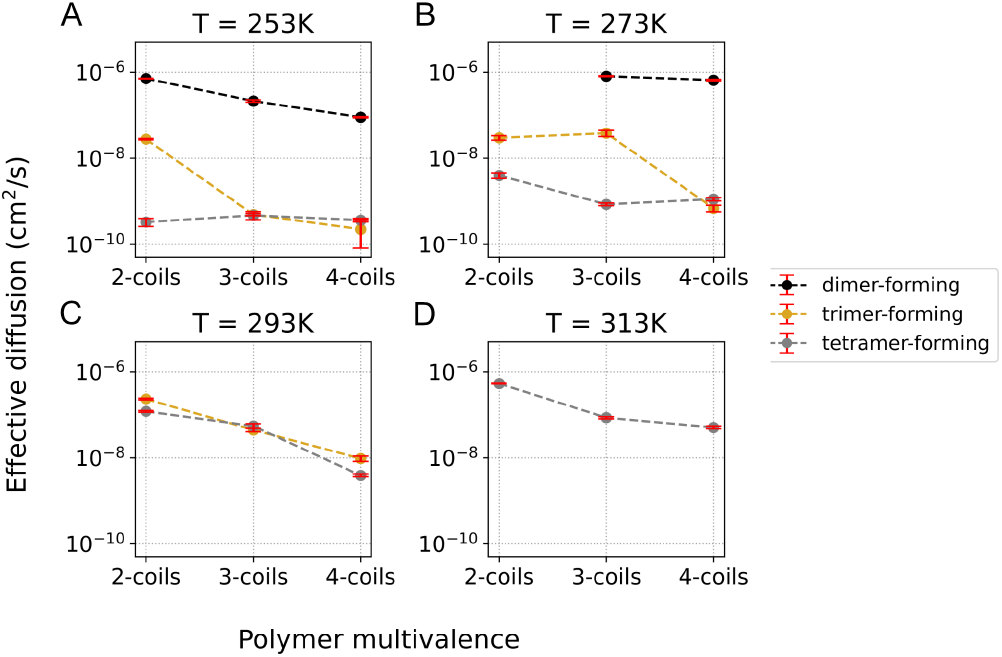
Multimer multivalency greatly impacts the dynamics of proteins within droplets. Plots of effective diffusion coefficients at (A) 253 K, (B) 273 K, (C) 293 K, and (D) 313 K. Data on y-axis is shown in log-scale, and only proteins that form LLPS droplets are plotted. Average effective diffusion coefficient (circles) and standard deviation (red error bars) are estimated via bootstrapping on MSD analyses from three replicate simulations for each protein.

## DISCUSSION

We report in this paper: (1) a novel CG framework for studying the phase separation of CC proteins (CC LLPS framework); (2) evidence that CC domains can be sole drivers of LLPS in physically motivated coarse-grained protein models; and (3) evidence that multimerization of stickers has a strong influence on LLPS propensity and protein diffusion in droplets.

Our findings that CC protein models can LLPS over a range of temperatures (Fig. 5) show that CC domains can be plausible LLPS-driving features of proteins. This suggests that for proteins like spd-5 (*C. elegans*) or pericentrin (*H. sapiens*), one or more of their CC domains could be contributing to their LLPS. Indeed, experimental studies with some phase-separating CC proteins also suggests that CC-domains can drive LLPS. For example, truncation studies of TAZ (*H. sapiens*) and Ede1 (*S. cerevisiae*) that lack their CC domains show that these mutants lose the ability to LLPS, compared to wild-type (22, 23). A paper studying McdB (cyanobacteria *S. elongatus* PCC 7942) performed similar truncation studies and found that a large central CC domain is necessary for its LLPS (26). Thus, there is indication of LLPS driven by CC domains in a wide range of organisms from cyanobacteria to mammals. Rios et al. (18) identified CC domains important for the LLPS of spd-5 through cross-linking mass spectrometry, and they show that a large fraction of spd-5–spd-5 interactions occur through CC domains. They also identified a key interaction between a PReM-containing and CM2-like domain in phosphorylated spd-5 (18), both of which are CCs. This interaction is similar to an interaction found in centrosomin (14), an essential centrosomal protein in fruit flies. Our findings that CC domains can be sufficient for LLPS, combined with experimental evidence of CC-specific interactions in centrosomal proteins, further supports CC domains as potentially sole or primary drivers of LLPS in some proteins.

Our simulations demonstrate the importance of multimeric multivalency on protein LLPS. The flexibility to alter polymeric and multimeric multivalency is a feature we designed in our simulation framework to allow us to easily study the contribution of each type of multivalence to CC driven LLPS. By directly changing the number of coil segments in a protein and how those segments multimerize, we show that multimerization of coil segments has a stronger effect on LLPS propensity (Figs. 4 and 5) than polymeric multivalency. Analogous experiments with real or designed proteins would be challenging to accomplish, and simulations provide the advantage of testing hypotheses for how proteins interact to drive LLPS prior to experimental investigations. Other simulation studies have shown how polymeric multivalence affects LLPS (7, 53, 55–57), but our results are some of the first to investigate multimeric multivalency. Recent experimental work with spd-5 (17, 18) and McdB (26) demonstrates the importance of multimerization on CC-driven LLPS, and our results in combination with experimental findings support a model for multimerization as a valid mechanism in modulating protein LLPS. Further understanding of how CC proteins LLPS will require consideration for how CC domains multimerize, not just how many CC domains might be in a protein.

It is notable that coil multimerization state has such a dramatic effect on LLPS propensity and effective diffusion considering the framework’s only moderate control over specific multimer formation, e.g. simulations designed to contain tetramers also contain dimers and trimers at equilibrium (Fig. S4). The polymeric multivalence-dependent decrease in diffusion is in line with other studies which show that increasing the number of stickers in a protein increases the viscosity of the resulting condensate (58, 59), with the caveat that we measured effective protein diffusion instead of droplet viscosity. A lack of precise multimer formation somewhat limits our efforts to establish a quantitative relationship phase behavior and multimerization state. The trends we observe, however—that higher multimerization states result in greater LLPS propensity and reduced protein dynamics—are still valid. At a fixed multimerization state, polymeric multivalency has less impact on LLPS propensity as determined by apparent melting temperature. For any multimer-forming coil it is unknown how many additional coil segments are needed, above a 2-coil protein, to start seeing a significant increase in melting temperature. Additional studies may more clearly show the difference that polymeric versus multimeric multivalency has on these CG CC systems.

In our systems we restricted all CC interactions to interchain interactions, where only coil segments of a specific type can interact (Fig. 1E). Intrachain interactions, however, will also likely modulate LLPS propensity of CC proteins similar to what has been seen for IDPs. Dignon et al. (60) demonstrated that single molecule properties of IDPs, such as the theta temperature, correlate well with phase behavior. Similarly, random-phase-approximation theoretical analysis of charged IDPs show that radii of gyration correlates strongly with phase behavior (61). These findings suggest that for IDPs, intrachain contacts can become the interchain contacts necessary to support droplet formation. The interplay between intra- and interchain interactions is also known to affect LLPS behavior. Rana et al. (62) showed by simulation that for proteins with arbitrary numbers of stickers that can undergo both inter- and intrachain interactions, it is the pattern of the stickers that predicts their LLPS propensity. We expect that sticker patterning will also be important for CC driven LLPS, once intrachain interactions are introduced.

There are ways our framework could be modified to more accurately reflect biologically relevant CC proteins, which might be worth future investigation. These improvements could include: adding parallel vs. antiparallel coil orientations like those seen in real CC domains (40, 41); adding amino acid specific interaction terms into our framework, similar to CG force fields developed to study IDPs and LLPS (7, 52, 63); having coil monomers exist as partially unfolded and requiring them to pay a free energy cost of folding during multimerization, which is how real CC domains are thought to behave (64–68); and changing the size and effective solvation of linkers, which is known to affect protein LLPS propensity (33, 57, 69). The model in its current implementation, however, is well-suited to test the hypothesis of CC-driven LLPS and can be easily adapted to explore models of CC driven LLPS.

Coiled-coil domains as drivers of LLPS represents a new way to think about mechanisms of protein phase separation. The most frequent way to view stickers is as individual residues in intrinsically disordered regions of proteins, despite the flexibility of the stickers-and-spacers framework to be applied to various levels of protein organization (27). This is likely because a majority of proteins identified in biomolecular condensates are intrinsically disordered (70), making IDP components good candidates to study as drivers of LLPS. Our results provide additional evidence that coiled-coil domains could be valid stickers and should be studied in their own right in driving protein phase separation. Our study, in conjunction with recent knowledge about how spd-5 is thought to interact (17, 18), also justifies studying CC domains as drivers in the phase separation of centrosomal proteins such as pericentrin. Existing data is insufficient to say if CC proteins are the primary domains responsible for LLPS of centrosomal and similar proteins, but these results suggest that future studies to identify driving domains should examine intrinsically disordered and CC domains alike.

## Supporting information

Supporting Material Document 2 - IDP Rg references

## SUPPORTING MATERIAL DESCRIPTION

Online Supporting Material for this document can be found on bioRxiv https://www.biorxiv.org/. Document S1 (PDF) contains supplemental methods, figures S1–S10, and tables S1–S4.

Document S2 (Excel file) contains a table of IDPs and associated radii of gyration, along with citations for the experimentally determined values, used in the validation of the linker segments.

## ACKNOWLEDGMENTS

We thank Dr. Roman Jerala and Maruša Ramšak for useful discussions about implementing coiled-coil interactions into framework. We also thank Dr. Chris Walker for his ideas during the early stages of framework development and in his insights in the validation of coil segment dimer dynamics. This work utilized the Alpine high performance computing resource at the University of Colorado Boulder. Alpine is jointly funded by the University of Colorado Boulder, the University of Colorado Anschutz, Colorado State University, and the National Science Foundation (award ACI-2201538). This work also used the Bridges-2 supercomputing system, which is supported by NSF award number ACI-1928147, at the Pittsburgh Supercomputing Center (PSC), from the Advanced Cyberinfrastructure Coordination Ecosystem: Services & Support (ACCESS) program, which is supported by National Science Foundation grants nos. 2138259, 2138286, 2138307, 2137603, and 2138296. This work was supported by the NIH Molecular Biophysics Training Program (T32GM065103, D.A.R.) and NSF MCB-1943488 (L.H.).

## AUTHOR CONTRIBUTIONS

Contributions according to CRediT Taxonomy:

**D. A. R**.: Methodology, software, investigation, visualization, writing. **L. E. H**.: Conceptualization, writing, supervision. **M. R. S**.: Conceptualization, writing, supervision, project administration.

## SUPPORTING CITATIONS

References (71–75) appear in the Supporting Material.

