## Supporting Material Document 1 for "Coiled-coil domains are sufficient to drive liquid-liquid phase separation of proteins in molecular models"

### SUPPLEMENTAL METHODS

#### S.I Structural parameters of CC LLPS framework

All CG beads are assigned a mass of 109.0 amu, which is approximately the average mass of the twenty proteinaceous amino acids. Bond stretching between all CG beads is treated with a harmonic potential with an equilibrium distance of 0.385 nm (to match the average distance between C$\alpha$ carbons) and a stiff force constant of 50000 kJ/mol/nm$^2$.

Pseudo-bond angles and pseudo-torsions were selected based off of $\phi$,$\psi$ angles from select human CC proteins including keratin (PDB: 3TNU), laminin (PDB: 1X8Y), vimentin (PDB: 3KLT), and a fragment of $\beta$-myosin (PDB: 2FXM) using `ProDy` v2.3.1 (1, 2). The average $\phi$,$\psi$ for the four CC proteins is (-64.088°, -41.260°). Applying the conversion formula from Tozzini et al. (3) results in pseudo-bond and pseudo-torsion values of 93 and 50 degrees, respectively. Pseudo-bond angles are treated with a harmonic potential, with an equilibrium angle of 93 degrees and a force constant of 50.0 kJ/mol/rad$^2$. Each pseudo-torsion is treated by a linear combination of two periodic cosine potentials. The first potential uses an equilibrium angle of -130.0 degrees (corresponding to a pseudo-torsion value of 50 degrees), a force constant of 20 kJ/mol, and multiplicity factor of 1. The second potential uses an equilibrium angle of -30.0 degrees, a force constant of 20 kJ/mol, and a multiplicity factor of 3. The above parameters are used exactly for the coil segment structure. The force constants were chosen because they result in coil segments that preserve helical shape but allow some small flexibility in the long axis of the coil so that the structure is not like a stiff rod. The linker segments use the exact angles as listed above, but with force constants for pseudo-bond angles and pseudo-torsions that are 100× weaker than the coils.

The number of neighbors excluded from nonbonded interactions, for any given bead, is 4. Therefore, explicit 1-4 and 1-5 nonbonded pair interactions are used in both the coil and linker segments, with a 12-6 Lennard-Jones potential. 1-4 pairs interact with $\epsilon$ = 2 kJ/mol and $\sigma$ = 0.453 nm. 1-5 pairs interact with $\epsilon$ = 2 kJ/mol and $\sigma$ = 0.561 nm. Linker segment beads use the same sigmas for the 1-4 and 1-5 pairs, but with 100x weaker $\epsilon$. 1-4 and 1-5 pair interactions do not cross between coil and linker segment beads.

#### S.II Interaction parameters of CC LLPS framework

We use three classes of bead types in the CC LLPS framework: coil-coil sticky beads, multimer-driving beads, and inert beads (diagram in Fig. 1D). Coil segment beads are organized to reproduce a heptad-repeat like pattern, similar to real coiled-coil domains. The heptad repeat, with amino acid positions labeled *'abcdefg'*, is a common feature seen in CC domains (4, 5). Residues in positions *'a'* and *'d'* are usually hydrophobic and drive the association of CC domains (4). Residues in positions *'e'* and *'g'* provide additional interactions between CC domains and are important for controlling multimerization state (5).

Coil-coil sticky beads are responsible for mediating the interactions between coil segments. These beads are placed in positions *'a'* and *'d'* of the heptad repeat (Fig. 1D) and are the primary drivers of coil segment association. The multimer-driving beads help control the type of multimer that coil segments can form. These beads are placed in positions *'e'* and *'g'* and flank the sticky beads (Fig. 1D). Interaction distances for coil-coil sticky beads (in positions *'a'*–*'a'*/*'d'*–*'d'* of the heptad repeat) and for multimer-driving beads (positions *'e'*–*'g'*) between associating coil segments are based off of distances between C-$\alpha$ carbons in similar positions from real CC proteins (described in Supporting Material section S.III, Fig. S1). This distances shown in Figure S1 served as starting points for optimization but do not represent the actual values used in the framework.

The remainder of a coil (positions *'b'*, *'c'*, and *'f'*) is occupied by inert beads, which are also the beads used to make the entirety of linker segments. These beads use relatively weak interaction strengths so they do not contribute significant protein-protein interactions. All of the bead interaction terms were parameterized to reflect the behavior of CC domains and disordered domains in solution, which allows us to use this framework with the solvent treated implicitly. GROMACS allows the use of a modified 12-6 Lennard-Jones potential:

$$V_{LJ}(r_{rj}) = \frac{C_{12,ij}}{r_{ij}^{12}} - \frac{C_{6,ij}}{r_{ij}^{6}}$$

where,

$$C_{12,ij} = 4\epsilon_{ij}\sigma_{ij}^{12}, \text{ and } C_{6,ij} = 4\epsilon_{ij}\sigma_{ij}^{6}$$

The use of the combined terms $C_{12,ij}$ and $C_{6,ij}$ allows us to set the attractive component of the Lennard-Jones potential equal to zero. We used this approach for the inert beads such that all interactions involving an inert bead have no attractive component of the potential but would retain the repulsive component. Table S2 lists all the interaction terms used in the framework for all multimeric states; the $C_{12,ij}$ and $C_{6,ij}$ for interactions between each of the bead types is also listed. No explicit Coulombic interaction terms were used in this study.

#### S.III Analysis of coiled-coil protein structures to select interaction parameter distances

We analyzed published crystal structures of CC proteins of various multimerization states to rationally choose starting $\sigma$ parameters for both the coil-coil sticky and multimer-driving beads. We selected 6 structures each of dimer-, trimer-, and tetramer-forming coiled-coil proteins. Only parallel CC proteins were selected for this analysis. We used DeepCoil V2 (6) to identify the location of the CC domain in each protein, as well as the starting *'a'* or *'d'* position residue of the first heptad. We calculated the distances between the C-$\alpha$ carbons of *'a'–'a'*, *'d'–'d'*, and *'e'–'g'* position residues in each heptad, for all heptads, between each coil and its nearest neighbors using `ProDy` v2.3.1 (1, 2). The average distances of all *'a'–'a'/'d'–'d'* and *'e'–'g'* distances in a protein were averaged, and then the average distances from each protein were averaged across all six proteins analyzed and are reported in Figure S1, showing the mean and standard deviation (error bars). Table S1 lists all the proteins used in this analysis, including the PDB codes, and the starting and ending amino acid indices of each CC domain relative to the numbering scheme in the respective PDB file.

#### S.IV Encoding interaction specificity between coil segments

We added only interchain attractive interactions in our CC proteins, and only between coil segments of the same ID (see Fig. 1E for a visual depiction). We did not add any attractive intrachain coil interactions in this study. We denote this as the *maximally specific* interaction regime, as it eliminates all possibility of intrachain interactions. To achieve this, for proteins with more than one coil segment, we made unique bead identifiers for each of the coil segments' coil-coil sticky beads. Coil segments with the same coil-coil bead ID interact using the interaction parameters specified above. Coil segments with different coil-coil bead IDs have no attractive interaction term specified between them (i.e. use the same interaction terms as the inert beads).

#### S.V Temperature replica exchange MD to assess structure of coarse-grained CC dimer

A CC dimer composed of two, 32-amino acid long coiled-coils was made using the CC Builder 2.0 web tool (7). The generated coil dimer was converted into a C-$\alpha$ CG representation with `ProDy` v2.3.1 (1, 2) and structural parameters from our framework were applied. We performed 1 $\mu$s temperature REMD simulations of the coil dimer, with 8 replicas between 277–411 K, and tested the effect of various coil-coil sticky bead interaction strengths on dimer stability. We tested $\epsilon$ values of 4.0, 5.0, 5.5, and 6.5 kJ/mol. Temperature replicas were first energy minimized, then equilibrated in the NVT ensemble for 150 ps with a time step of 10 fs. Temperature REMD simulations were then done in the NVT ensemble for 1 $\mu$s and a time step of 10 fs, with replica exchanges attempted every 200 steps. We determined the medoid structure from each replica's trajectory (which is continuous in temperature) using the DBSCAN clustering method. The backbone RMSD of coils from each replica was calculated using the respective medoid structure as the reference. We implemented our RMSD analysis to account for symmetry in the coil dimer structure so that parallel and antiparallel dimers are treated equally. We reconstructed the reference RMSD distribution (8) (Fig. S2A) and fit Gaussian kernel density estimates to the reference and all CG RMSD distributions using `scipy` v1.9.1. RMSD distributions from each tested $\epsilon$ value are shown from simulations at 293 (Fig. S2B) and 310 K (Fig. S2C). We calculated the Kullback-Leibler between each CG kernel to the reference kernel, and used these values to determine the similarity between our data to the reference.

#### S.VI Single molecule MD of intrinsically disordered proteins to parameterize linker segments

We validated the design of our linker segments and show that we can reproduce experimental radii of gyration ($R_g$) of real IDPs. We designed the linker segments so that they would be disordered, not contribute protein-protein interactions, and interact equally well with (implicit) solvent and protein components. Our goal with these design constraints was to create a linker segment whose behavior is similar to the dynamics of real IDPs. We selected 23 different IDPs from Dignon et al. (9) and Tesei et al. (10), covering a range of 24 to 196 amino acids (CG beads), to turn into linker segments for single molecule simulation. Protein names, other identifiers and sequences (where applicable), and experimentally determined radii of gyration ($R_g$) with error estimates plus the temperature they were determined at for all 23 proteins are reported in Supporting Material, Document 2. This document also lists the original publication which reported the $R_g$ for each protein. The 23 proteins were turned into linkers using the `PeptideBuilder` strategy (Supporting Material, section S.VIII), energy minimized, and then equilibrated in the NVT ensemble for 250 ps with a time step of 10 fs. Production NVT simulations were then performed for 300 ns with a time step of 10 fs. The temperature of equilibration and production for each linker reflected the experimental temperature at which the $R_g$ for the corresponding IDP was determined. Simulations of each linker were done in triplicate. The first 50 ns of production was discarded for equilibration, and the average $R_g$ was calculated from the remaining trajectory. Mean $R_g$ and standard deviation are reported. $R_g$ was calculated with `mdtraj` v1.9.7 and the correlation coefficient was calculated with `scipy` v1.9.1.

#### S.VII MD to assess multimerization capacity of coil segments

We designed three 3-coil-2-linker CC proteins using the `PeptideBuilder` strategy (Supporting Material, section S.VIII). We specified the coils either as dimer-forming, trimer-forming, or tetramer-forming. We ran single molecule simulations for all three of these proteins. We energy minimized, equilibrated in the NVT ensemble at 298.15 K for 100 ns with a timestep of 25 fs, and then did production simulation in the NVT ensemble at 298.15 K for 1 $\mu$s with a timestep of 25 fs. The final configuration of each protein was then used to pack a box with 8 copies of the protein using `packmol` v20.11.1 (11), into a 20x20x20 nm box with a packing tolerance of 1 nm. At this point, there were three separate packed boxes, one filled with dimer-forming coil proteins, one with trimer-forming coil proteins, and one with tetramer-forming coil proteins. We then ran triplicate MD simulations on each packed box. Packed boxes were energy minimized, equilibrated in the NVT ensemble at 298.15 K for 100 ns with a timestep of 25 fs, and then simulated in the NVT ensemble at 298.15 K for 2 $\mu$s with a timestep of 25 fs. We quantified the number of multimers in each replicate simulation using a custom python script. Our analysis method determines coil multimers in desired frames of the simulation by calculating the distances between all coil segments' center of mass and finding neighbors within a 1.3 nm cutoff. The output from this analysis is the number of each type of multimer (monomer through hexamer) per frame. We then average the number of each multimer type across all analyzed frames and normalize to the total number of coil segments in the simulation (in this case, 24) to determine the fraction of each multimer in a simulation. These values are then averaged across the three simulation replicates. The first 100 ns of each packed box simulation was discarded before analysis.

Multiple simulations starting from the packed box but with varied interaction parameters were performed to select the appropriate parameters for the coil-coil sticky and multimer-driving beads. Starting $\epsilon$ values for trimer- and tetramer-forming coils were chosen from dimer-forming coils, and starting $\sigma$ values were based on the distances calculated in Figure S1. Interaction parameters were varied until the desired multimer populations for each multimer type was achieved (Fig. S4).

#### S.VIII Generating CG coiled-coil proteins

Whole coarse-grained CC proteins in our framework are constructed using the python package `PeptideBuilder` v1.1.0 (12) and are converted into C-alpha coarse-grained representations using the python package `ProDy` v2.3.1 (1, 2). All parts of the protein, including the coil and linker segments, are initially assembled in helical geometry with phi/psi angles of (-64.088°, -41.260°; see Supporting Material section S.III for an explanation of these values). `PeptideBuilder` requires an amino acid sequence to construct a protein, and so we designed fake sequences to first build the proteins. The choice of these fake sequences is arbitrary and all sequences will make the same initial backbone. We have the ability to choose where along the coil segment the first heptad starts, but for all proteins in this sutdy we begin the first heptad at the first bead in a coil segment. Helper scripts for generating 3D structures along with topology files with proper parameters can be found on the GitHub repository (`https://github.com/dora1300/cc_llps_framework`). Throughout this study, the lengths (no. of beads) of coil and linker segments is fixed to 32 and 25 beads, respectively. These values represent the average median length of coiled-coil domains and intrinsically disordered regions from centrosomal proteins CDK5RAP2, spd-5, and centrosomin. All proteins generated this way require subsequent MD to relax the linker segments out of the helical geometry and into their equilibrium pseudo-angles/torsions.

#### SUPPLEMENTAL FIGURES

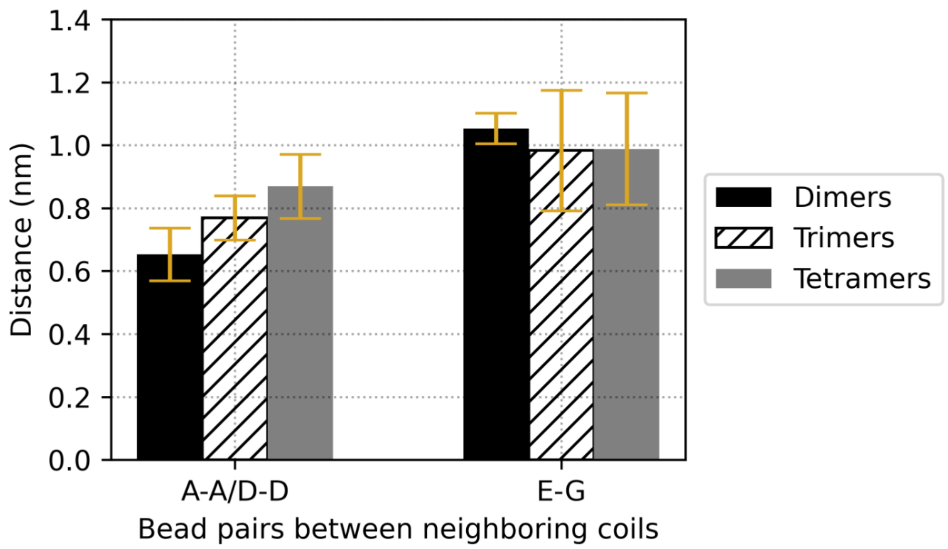

Figure S1: Multimer-dependent distances between residues in real CC proteins. Data are mean ± standard deviation ($N$=6 proteins for each multimer type) of distances between C-$\alpha$ carbons of neighboring CC domains from crystal structures of CC proteins for each of the heptad positions listed. The structures analyzed to generate these data are listed in Table S1.

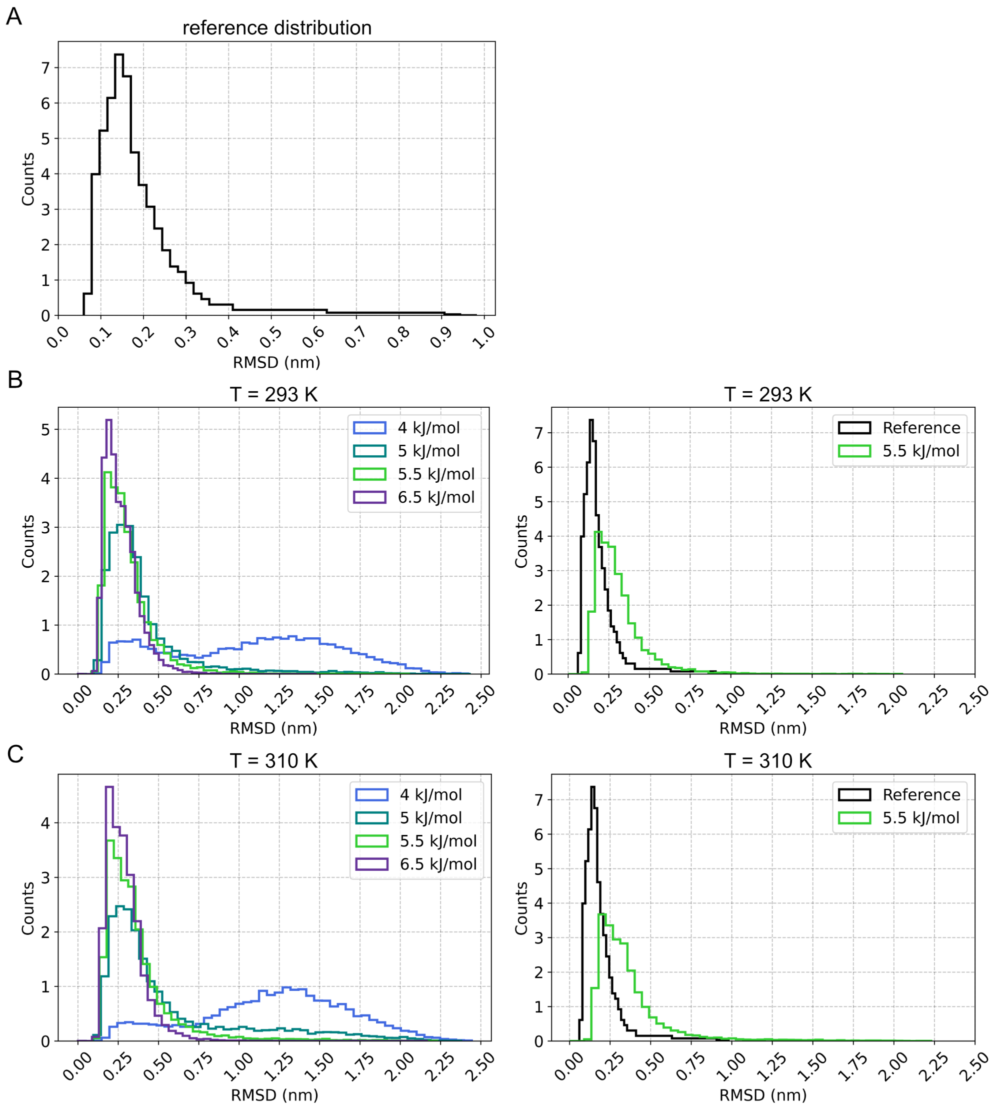

Figure S2: Coarse-grained coil dimers behave like atomistic coil dimers. (A) Plot of reconstructed normalized reference backbone RMSD, representing an approximated average of distributions from atomistic coiled-coil dimer simulations across 64 replicas spanning 298 to 419 K (8). (B) Plot of RMSD counts of a CG coil dimer from REMD simulations at different coil-interaction strength at 298 K (left). RMSD is calculated with reference to the medoid structure from each interaction strength's simulation. The panel on the right compares the chosen optimal interaction strength RMSD distribution (5.5 kJ/mol) superimposed against the reference. (C) The same type of data as presented in *(B)* but at 310 K.

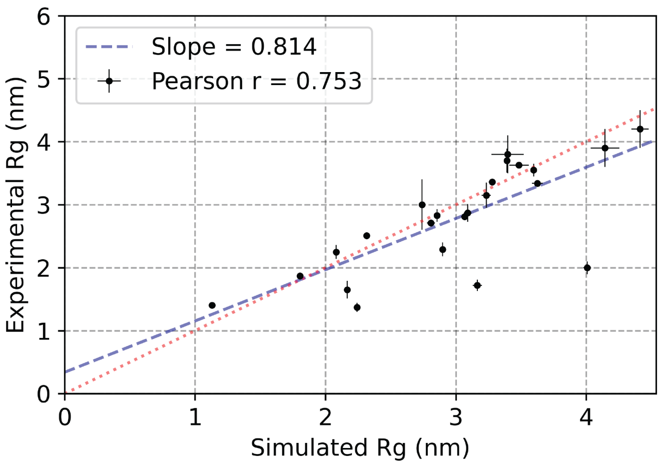

Figure S3: Simulated linkers reproduce experimental $R_g$ of disordered proteins. Data points are the average $R_g$ from three replicate simulations of one type of linker, plotted against the experimental $R_g$ for the corresponding IDP. Error bars are standard deviation from simulation (in $x$-direction) and from experiment (in $y$-dimension, see Supporting Document 2). Blue dashed line is the line of best fit for the data, and red dotted line is the parity line.

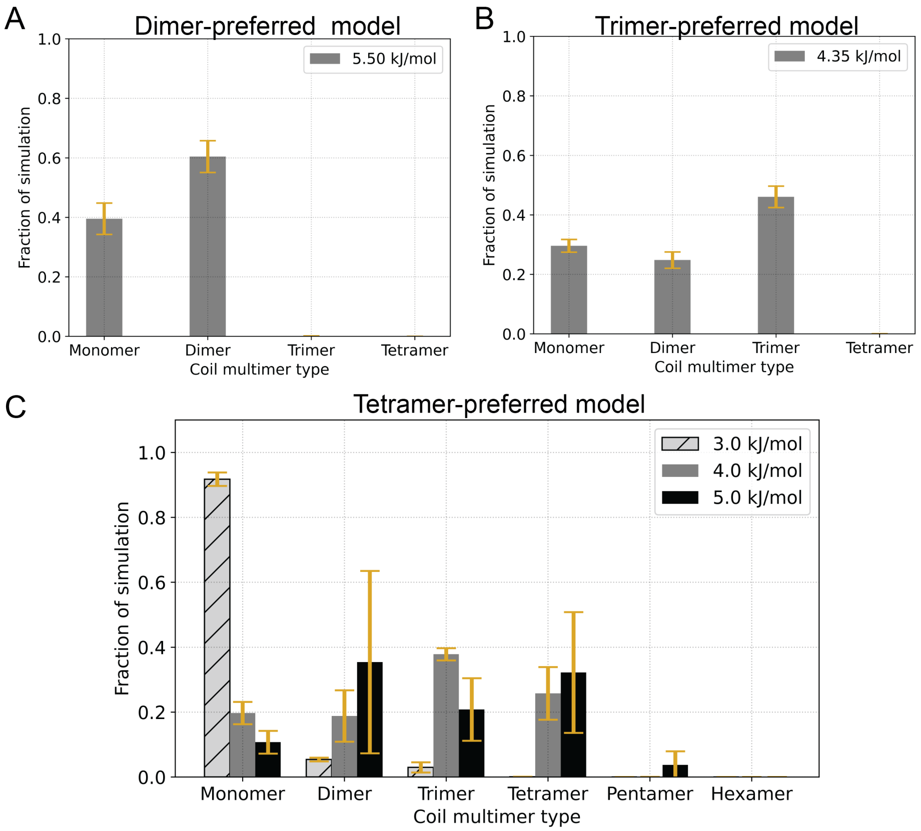

Figure S4: Multimerization of coil segments is qualitatively controllable. For all plots, data are mean ± standard deviation from three replicate simulations. (A) Relative probability of multimer species from simulations of dimer-forming coils. Value in legend is interaction value for the dimer specific coil-coil sticky beads. (B) Relative probability of multimer species from simulations of trimer-forming coils. Value in legend is interaction value for the trimer specific coil-coil sticky beads. (C) Relative probability of multimer species from simulations of tetramer-forming coils. Values in legend compare the effect that tetramer specific coil-coil sticky bead has on multimer formation.

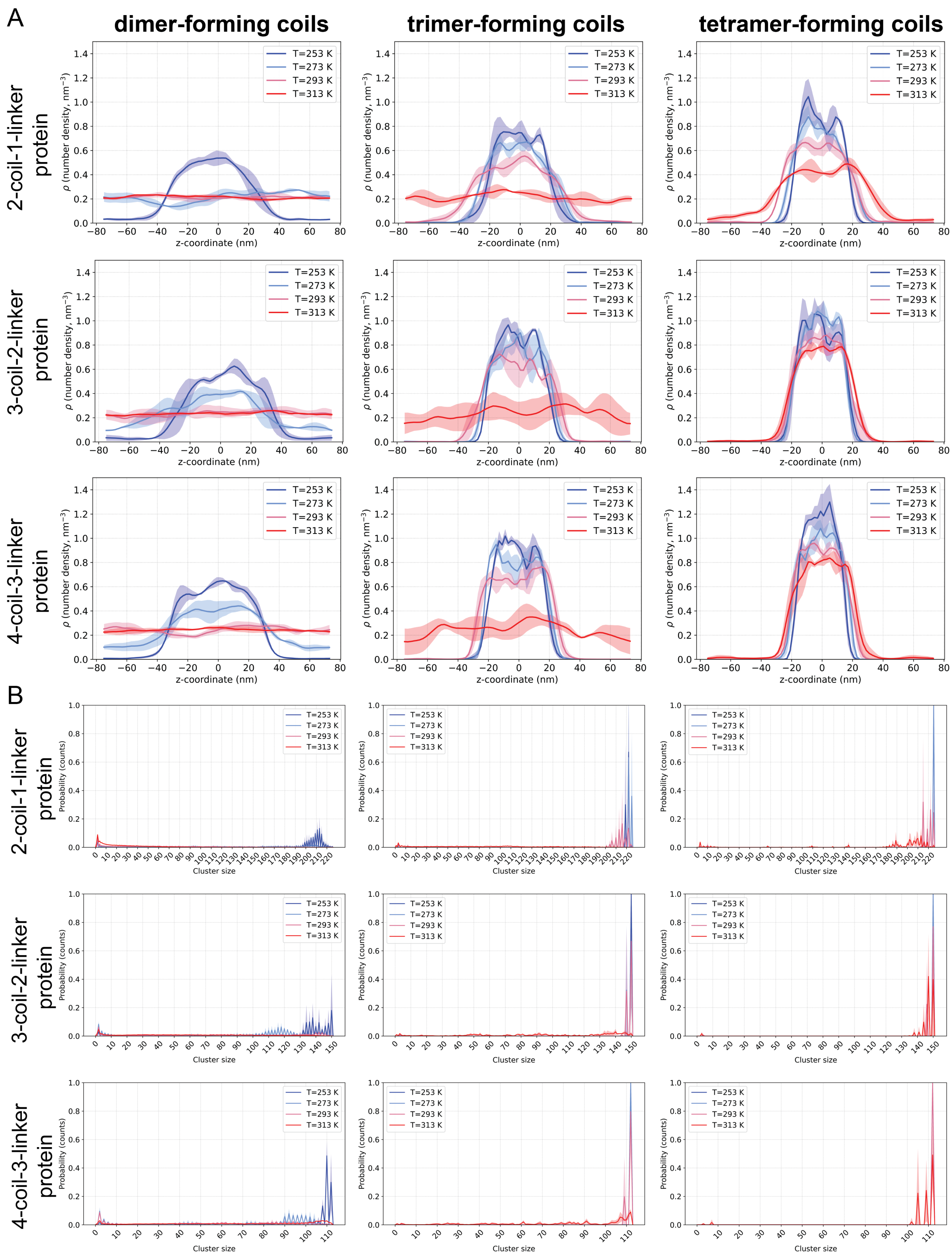

Figure S5: Density profile and molecular cluster size analyses from equilibrated slab simulations of (A) 2-coil-1-linker CC proteins, (B) 3-coil-2-linker CC proteins, and (C) 4-coil-3-linker CC proteins. Columns are differentiated by the coil-multimer type (dimer:left, trimer:middle, tetramer:right). In all plots, solid lines represent mean, and shaded regions are standard deviation, from three replicate simulations.

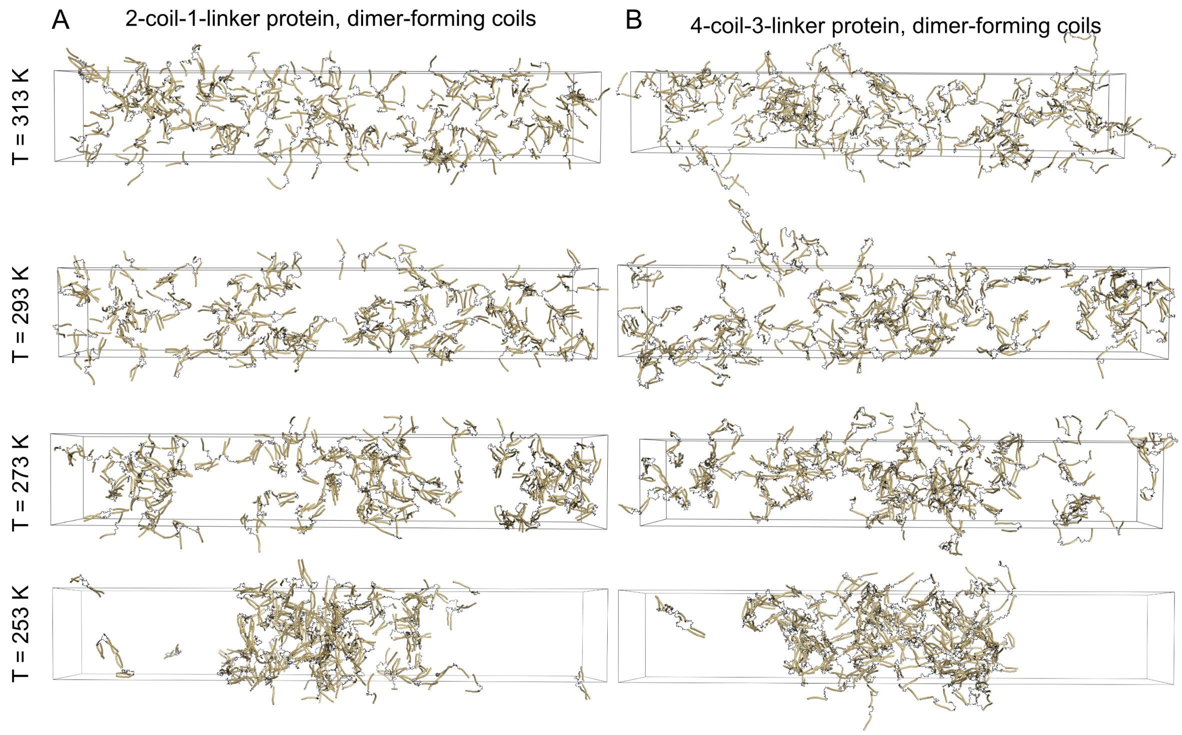

Figure S6: LLPS behavior is dependent on temperature for dimer-forming proteins. Snapshots (at 20 $\mu$s) from slab simulations from one replicate set of (A) 2-coil and (B) 4-coil dimer-forming proteins. Images are organized similarly to Figure 3 but without corresponding binodal plots (which are presented in Fig. 5). Snapshots at 253 K for both (A) and (B) are the same as in Figure 2. Proteins are made whole for visualization, but would wrap through the box boundaries during actual simulation. Visualizations produced using open-source PyMOL v.2.5.0.

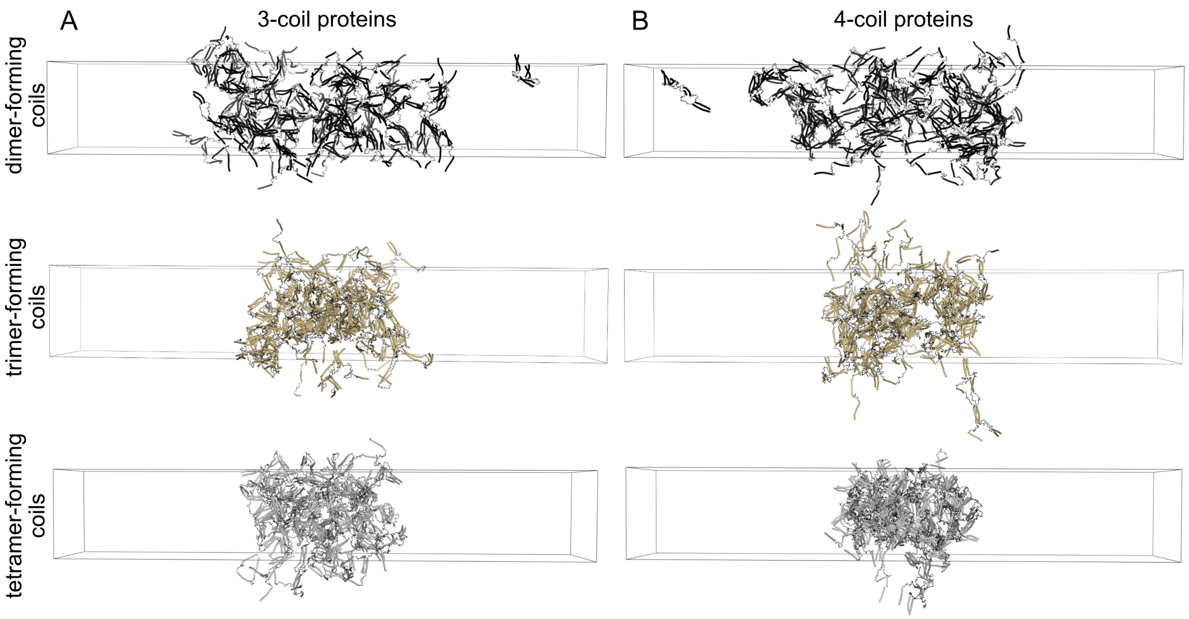

Figure S7: The size of slabs is dependent on multimer forming state. Representative snapshots (at 20 $\mu$s) from slabs of (A) 3-coil proteins and (B) 4-coil proteins at 253 K, for proteins with dimer-forming (top row, black), trimer-forming (middle row, gold), and tetramer-forming (bottom row, silver) coils. Snapshots from dimer-forming coil simulations (top row) are the same as in Figure 2. Proteins are colored the same as in Figure 4. Proteins are made whole for visualization but would wrap through the box boundaries during simulation. Visualizations produced using open-source PyMOL v2.5.0

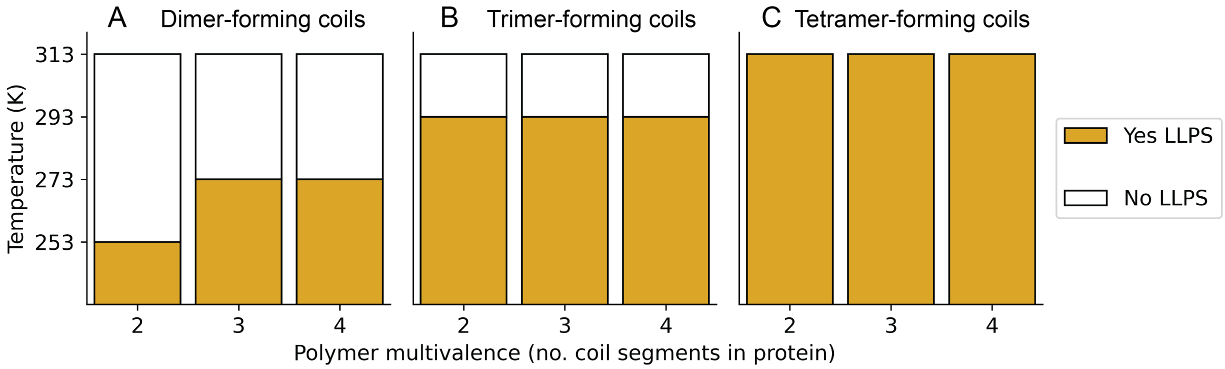

Figure S8: Polymeric multivalency only weakly impacts LLPS propensity. Phase diagrams comparing the effect of 2-, 3-, and 4-coils on LLPS propensity for (A) dimer-forming coils, (B) trimer-forming coils, and (C) tetramer-forming coils. Bars corresponding to phase behavior are determined qualitatively from density profiles and molecular cluster analyses of simulations (Fig. S5).

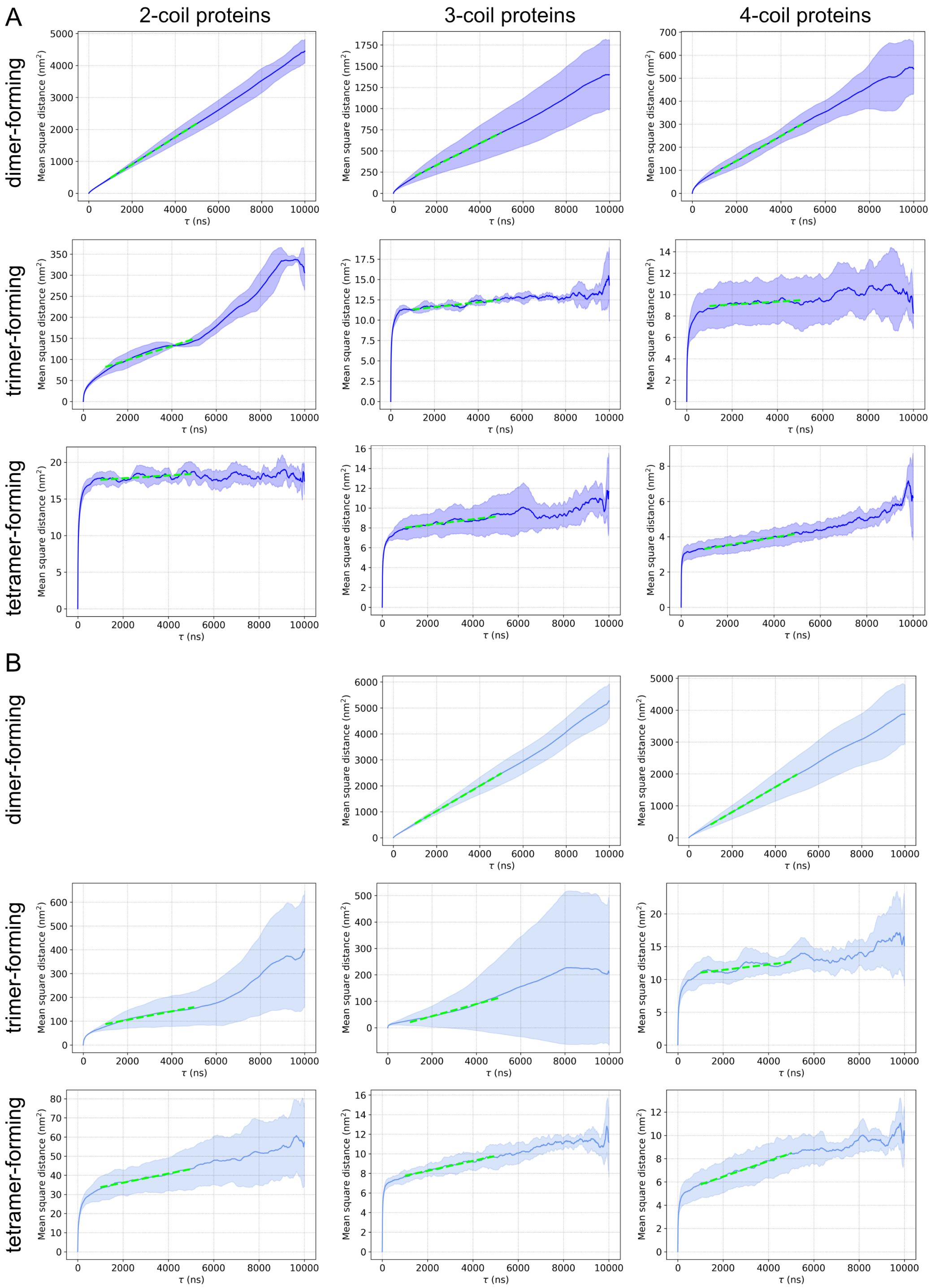

Figure S9: Estimation of diffusion coefficients from MSD analysis at 253 and 273 K. Individual plots show both the average MSD of individual proteins for each protein along with a line representing the linear fit from bootstrap analysis (green dashed line) used to estimate the effective diffusion coefficient at (A) 253 K and (B) 273 K. See Methods for additional details about bootstrap analysis. Average MSD (solid line) and standard deviation (shaded regions) are calculated from three simulation replicates. Only data for proteins that form LLPS droplets are presented.

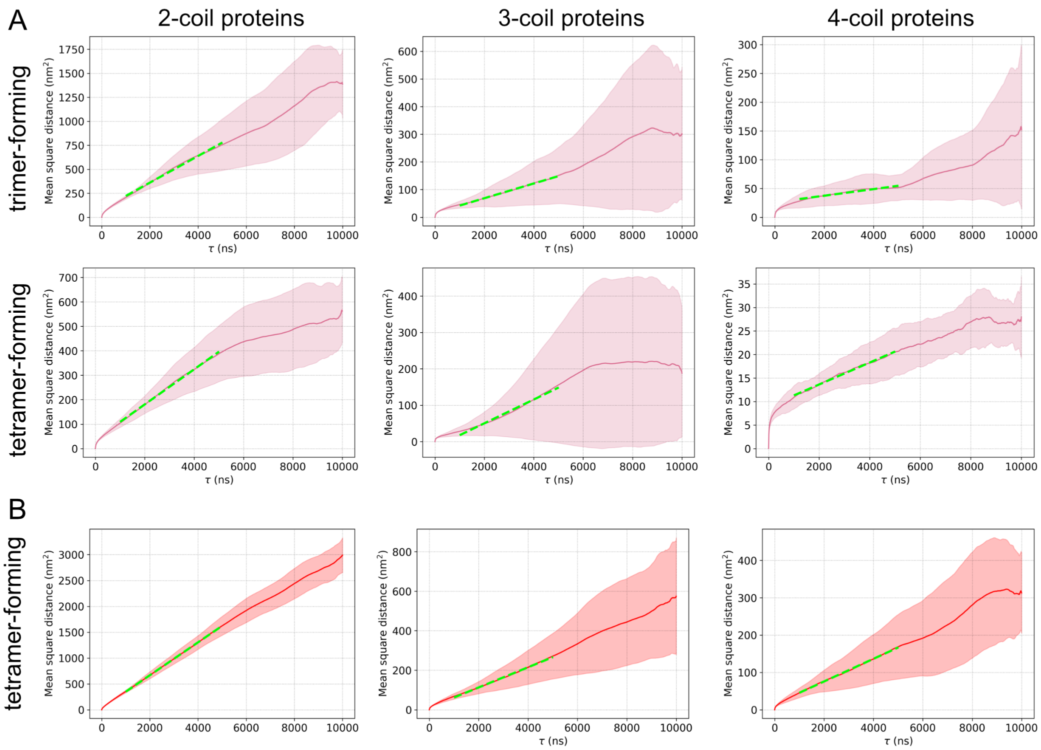

Figure S10: Estimation of diffusion coefficients from MSD analysis at 293 and 313 K. Individual plots show both the average MSD of individual proteins for each protein along with a line representing the linear fit from bootstrap analysis (green dashed line) used to estimate the effective diffusion coefficient at (A) 293 K and (B) 313 K. See Methods for additional details about bootstrap analysis. Average MSD (solid line) and standard deviation (shaded regions) are calculated from three simulation replicates. Only data for proteins that form LLPS droplets are presented.

#### SUPPLEMENTAL TABLES

| Multimer type | PDB code | Starting register | Starting index | Ending index |
|---|---|---|---|---|
| **Dimer** | 4DZM | '*a*' | 3 | 31 |
| | 2FXM | '*a*' | 856 | 957 |
| | 2D3E | '*a*' | 155 | 270 |
| | 1D7m | '*a*' | 248 | 339 |
| | 2B9C | '*a*' | Chain A: 113<br>Chain B: 1113 | Chain A: 225<br>Chain B: 1225 |
| | 1GK4 | '*a*' | 337 | 404 |
| | 1C1G | '*a*' | 8<br>Chain B: offset by 284 | 274 |
| **Trimer** | 4DZK | '*a*' | 3 | 30 |
| | 4DZL | '*a*' | 3 | 30 |
| | 1AA0 | '*a*' | 442 | 454 |
| | 1KFN | '*a*' | 6 | 50 |
| | 4DZN | '*a*' | 3 | 30 |
| | 6FIA | '*a*' | 73 | 146 |
| **Tetramer** | 6XY1 | '*d*' | 3 | 29 |
| | 6XXZ | '*d*' | 3 | 29 |
| | 1NHL | '*d*' | 32 | 78 |
| | 1EZJ | '*d*' | 75 | 92 |
| | 6XY0 | '*d*' | 3 | 28 |
| | 3DZO | '*d*' | 317 | 370 |

Table S1: PDB IDs, and starting and ending indices used in the analysis of C-$\alpha$ distances between multimerized coiled-coil proteins.

| | Bead pairs | Bead class | $\epsilon$ (kJ/mol) | $\sigma$ (nm) | $r_{min}$ (nm) | $C_6$ (attractive) | $C_{12}$ (repulsive) |
|---|---|---|---|---|---|---|---|
| **Dimer beads** | | | | | | | |
| | $A_{dim}$–$A_{dim}$ | coil-coil sticky | 5.5 | 0.570 | 0.640 | 7.559142E-01 | 2.597302E-02 |
| | Di–Di | multimer-driving | 3.0 | 0.891 | 1.00 | 6.00 | 3.00 |
| **Trimer beads** | | | | | | | |
| | $A_{tri}$–$A_{tri}$ | coil-coil sticky | 4.35 | 0.570 | 0.640 | 5.978594E-01 | 2.054229E-02 |
| | Tri–Tri | multimer-driving | 2.0 | 0.802 | 0.90 | 2.125764 | 5.648591E-01 |
| **Tetramer beads** | | | | | | | |
| | $A_{tet}$–$A_{tet}$ | coil-coil sticky | 4.00 | 0.624 | 0.700 | 9.411920E-01 | 5.536515E-02 |
| | Tet–Tet | multimer-driving | 2.0 | 0.713 | 0.80 | 1.048576 | 1.374390E-01 |
| **Backbone and other beads** | | | | | | | |
| | B–B | inert backbone | 2.00 | 0.453 | 0.508 | 0.00 | 5.974044E-04 |
| | B–$A_M$ | inert | 2.00 | 0.453 | 0.508 | 0.00 | 5.974044E-04 |
| | B–*Multi* | inert | 2.00 | 0.453 | 0.508 | 0.00 | 5.974044E-04 |

Table S2: Interaction terms between bead types in CC LLPS framework. $A_M$ refers to any of the multimer-specific coil-coil sticky beads, and *Multi* refers to any of the multimer-driving beads.

| Interaction strength (kJ/mol) | Temperature | KL divergence (nats) |
|---|---|---|
| 4.0 | 293 | 2.38 |
| 5.0 | 293 | 0.96 |
| 5.5 | 293 | 0.90 |
| 6.5 | 293 | 1.06 |
| 4.0 | 310 | 3.33 |
| 5.0 | 310 | 1.11 |
| 5.5 | 310 | 1.01 |
| 6.5 | 310 | 1.34 |

Table S3: Kullback-Leibler divergences of backbone RMSD distributions between CG and atomistic simulations of a coiled-coil dimer. Divergences were calculated using Gaussian kernel density estimates shown in Figure S2.

| Effective diffusion coefficients ($\times 10^{-9}$) ($cm^2/s$) | | | |
|---|---|---|---|
| **253 K** | | | |
| | 2-coil | 3-coil | 4-coil |
| dimer-forming | 712.0 ± 12.06 | 213.5 ± 11.84 | 89.47 ± 2.282 |
| trimer-forming | 27.71 ± 0.8990 | 0.4895 ± 0.0303 | 0.2228 ± 0.1420 |
| tetramer-forming | 0.3270 ± 0.0700 | 0.4685 ± 0.1011 | 0.3632 ± 0.0336 |
| **273 K** | | | |
| | 2-coil | 3-coil | 4-coil |
| dimer-forming | | 807.9 ± 18.82 | 653.8 ± 26.34 |
| trimer-forming | 29.73 ± 3.542 | 38.16 ± 6.146 | 0.6846 ± 0.1180 |
| tetramer-forming | 3.971 ± 0.5899 | 0.8441 ± 0.0595 | 1.119 ± 0.0955 |
| **293 K** | | | |
| | 2-coil | 3-coil | 4-coil |
| trimer-forming | 231.3 ± 11.42 | 44.40 ± 4.515 | 9.565 ± 1.344 |
| tetramer-forming | 119.5 ± 5.303 | 54.48 ± 6.782 | 3.881 ± 0.2553 |
| **313 K** | | | |
| | 2-coil | 3-coil | 4-coil |
| tetramer-forming | 530.2 ± 7.883 | 85.12 ± 4.983 | 50.54 ± 3.773 |

Table S4: Effective diffusion coefficients of proteins that undergo LLPS. All coefficients have units of $1 \times 10^{-9}$ $cm^2/s$.
